# Network-aware mutation clustering of cancer

**DOI:** 10.1101/432872

**Authors:** Swetansu Pattnaik, Catherine Vacher, Hong Ching Lee, Warren Kaplan, David M. Thomas, Jianmin Wu, Mark Pinese

**Affiliations:** The Kinghorn Cancer Centre and Cancer Division, Garvan Institute of Medical Research, 370 Victoria St, Darlinghurst, NSW, Australia.; Kinghorn Centre for Clinical Genomics, Garvan Institute of Medical Research, 370 Victoria St, Darlinghurst, NSW, Australia.; St Vincent’s Clinical School, Faculty of Medicine, UNSW Sydney, NSW, Australia.; Key laboratory of Carcinogenesis and Translational Research (Ministry of Education/Beijing), Center for Cancer Bioinformatics, Peking University Cancer Hospital & Institute.

## Abstract

The grouping of cancers across tissue boundaries is central to precision oncology, but remains a difficult problem. Here we present EPICC (Experimental Protein Interaction Clustering of Cancer), a novel technique to cluster cancer patients based on DNA mutation profile, that leverages knowledge of protein-protein interactions to reduce noise and amplify biological signal. We applied EPICC to data from The Cancer Genome Atlas (TCGA), and both recapitulated known cancer clusterings, and identified new cross-tissue cancer groups that may indicate novel cancer molecular subtypes. Investigation of EPICC clusters revealed new protein modules which were recurrently mutated across cancers, and indicate new avenues for research into cancer biology. EPICC leveraged the Vodafone DreamLab citizen science platform, and we provide our results as a resource for researchers to investigate the role of protein modules in cancer.

## Introduction

The grouping of cancers by molecular subtype, independent of site of origin, underpins precision oncology. This paradigm is transforming cancer treatment, but is critically dependent on the identification of molecular cancer subtypes that accurately stratify patients by clinical course and response to therapy.

Initial global approaches to defining cancer subtypes relied on transcriptome measurements (van’t Veer *et al.*, 2002). Although the transcriptome provides a holistic readout of cell state that is highly predictive of biological activity (Ray *et al.*, 2014), the lability of the RNA analyte limits its application in the clinic. This limitation has led to efforts to use more stable analytes to identify cancer subtypes, most notably somatic DNA changes.

Unfortunately, the analysis of DNA mutations suffers from a multiplicity issue: many different mutations, potentially affecting different genes, can have a similar biological effect. An example is the PI3K-AKT-mTOR pathway, which can be variously activated in cancer by gain of function mutations in *PIK3CA*, *AKT*, or mTOR, or loss of function changes in *PTEN*, *TSC1/2*, or *LKB1* (Janku *et al.*, 2018). Cancers sharing these various mutations should be logically grouped by their common dysregulation of mTOR, yet this similarity is obscured by mutations being spread across the many genes involved.

This challenge has previously been addressed by pathway analysis, in which genes are grouped into logical pathways, and cancers compared by their mutational profile at the pathway level (Sanchez-Vega *et al.*, 2018). These techniques are powerful but critically dependent on accurate knowledge of pathways and their components. As the majority of the human proteome remains unannotated for function (Mi *et al.*, 2017) this dependence on manually-defined pathways is a serious limitation.

Global databases of protein-protein interactions (PPIs) offer a solution. Proteins do not act in isolation, but rather interact, often through physical contact. These patterns of interaction can be determined experimentally, and define protein complexes and linked function in a manner that is not dependent on human annotation. The utilisation of PPIs to identify clinically relevant protein interaction subnetworks in the last decade has established networks science as an effective approach to interrogate disease biology (Li *et al.*, 2017; Ivanov *et al.*, 2018; Vinayagam *et al.*, 2011; Barabási *et al.*, 2011). However, these approaches have heavily relied on knowledge of sets of cancer driver genes and the directionality of the network architecture, and are limited in facilitating discovery of novel subnetworks. A need remains for an approach to cluster cancer patients by mutation profile, and discover protein modules driving the clustering, using a minimum of expert knowledge.

To address this need we have developed EPICC (Experimental Protein Interaction Clustering of Cancer), a novel technique to cluster cancer patients by mutational profile. EPICC combines the sensitivity of pathway-level analysis with the unbiased knowledge encoded in experimental protein-protein interaction networks, to cluster cancers by both known and novel biological patterns. We implemented EPICC in the Vodafone DreamLab mobile computation platform as Projects Decode and Genetic Profile, and applied it to mutation data from The Cancer Genome Atlas (TCGA), to reveal both known and novel patient clusterings and mutation patterns. EPICC provides a novel lens to analyse and group cancer mutations, and we provide our summary results as a resource for cancer researchers worldwide.

## Results

### EPICC: network-aware cancer clustering

EPICC is based on the intuition that protein-protein interactions are informative of shared function: proteins which physically interact are likely to be functionally linked also. EPICC applies this idea to increase the sensitivity of clustering cancers by the presence of shared mutations, by considering not only shared mutations in a single gene, but also shared mutations in gene neighbourhoods defined by the protein-protein interaction network.

At the core of EPICC is a metric that scores the similarity between two cancers by comparing their mutation profiles in the context of protein-protein interaction networks (Figure 1). The EPICC score between cancers A and B is defined as the number of protein-protein interactions between proteins mutated in A and proteins mutated in B. The key advantage of this score over other approaches is that it can detect similarity between patients even in the absence of common mutated proteins or any protein annotations.

**Figure 1.**
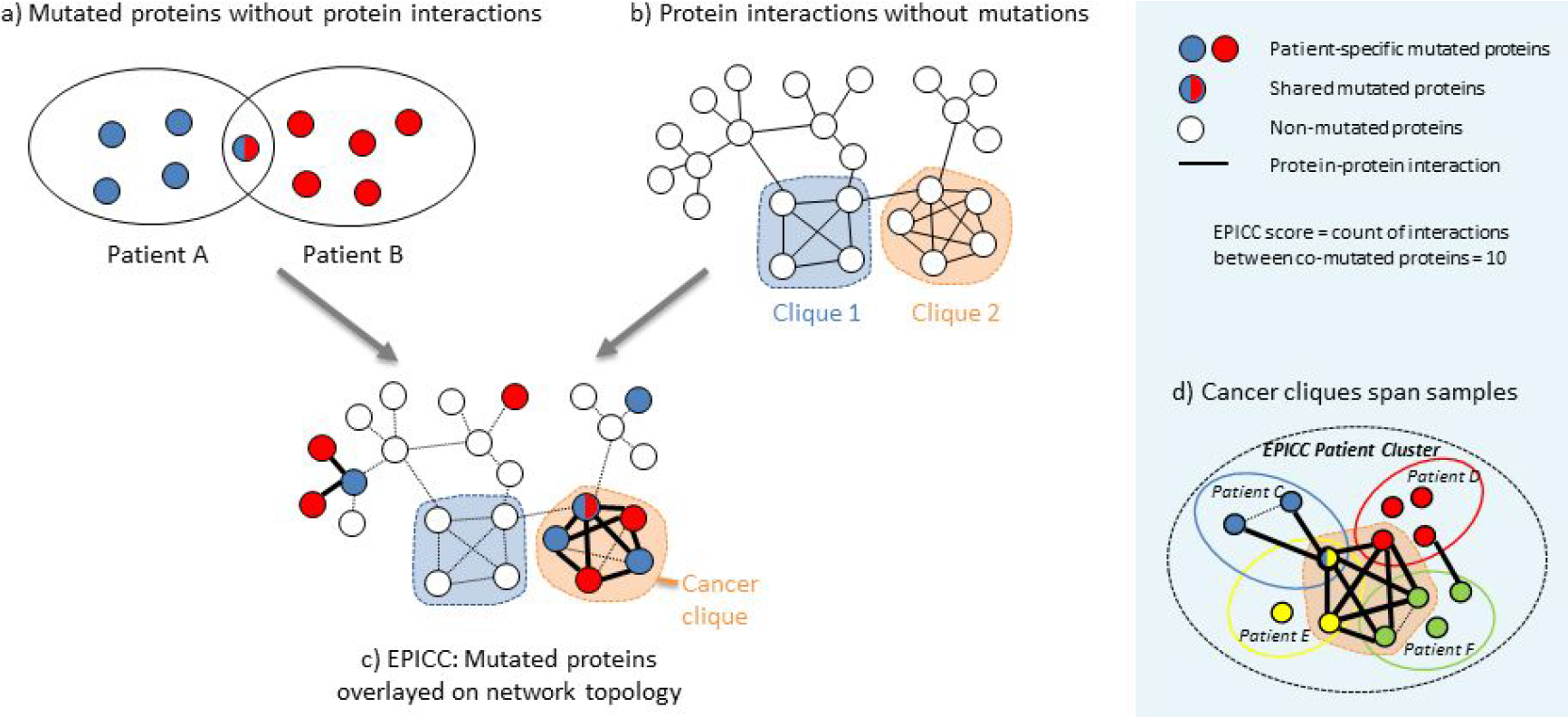
The EPICC concept. Comparing patients by the presence of overlapping mutated genes (a) is a straightforward approach, but does not leverage knowledge of protein-protein interactions. Conversely, the analysis of protein-protein interaction networks without mutation data (b) can identify fully-connected groups of proteins (cliques) which may form a biological unit, but cannot determine the importance of these cliques in cancer. EPICC combines these two approaches, by overlaying mutation data on to protein interaction networks (c). This enables the comparison of patient mutation profiles in a robust and network-aware manner, and also the identification of cancer cliques which may be important to cancer biology. Cancer cliques can span multiple patients (d), permitting the identification of consistently mutated protein modules even when the particular proteins mutated vary. Proteins are represented by circles coloured by the patients bearing the mutation. Protein-protein interactions are depicted by lines between proteins. In (c) and (d), interactions contributing to the EPICC score are shown as bold lines.

### EPICC clusters reveal known and new cancer groupings

We applied EPICC to mutation data from eight cancers profiled in The Cancer Genome Atlas project (TCGA), totalling 3,750 patients. 141 patient clusters were identified, with a wide distribution of sizes (Figure 2a). Although some clusters were highly specific to a certain cancer primary site, we observed a surprisingly broad representation of primary types within each cluster (Figure 2b). No clusters were associated with gender or race after controlling for primary site (Figure 2c).

**Figure 2:**
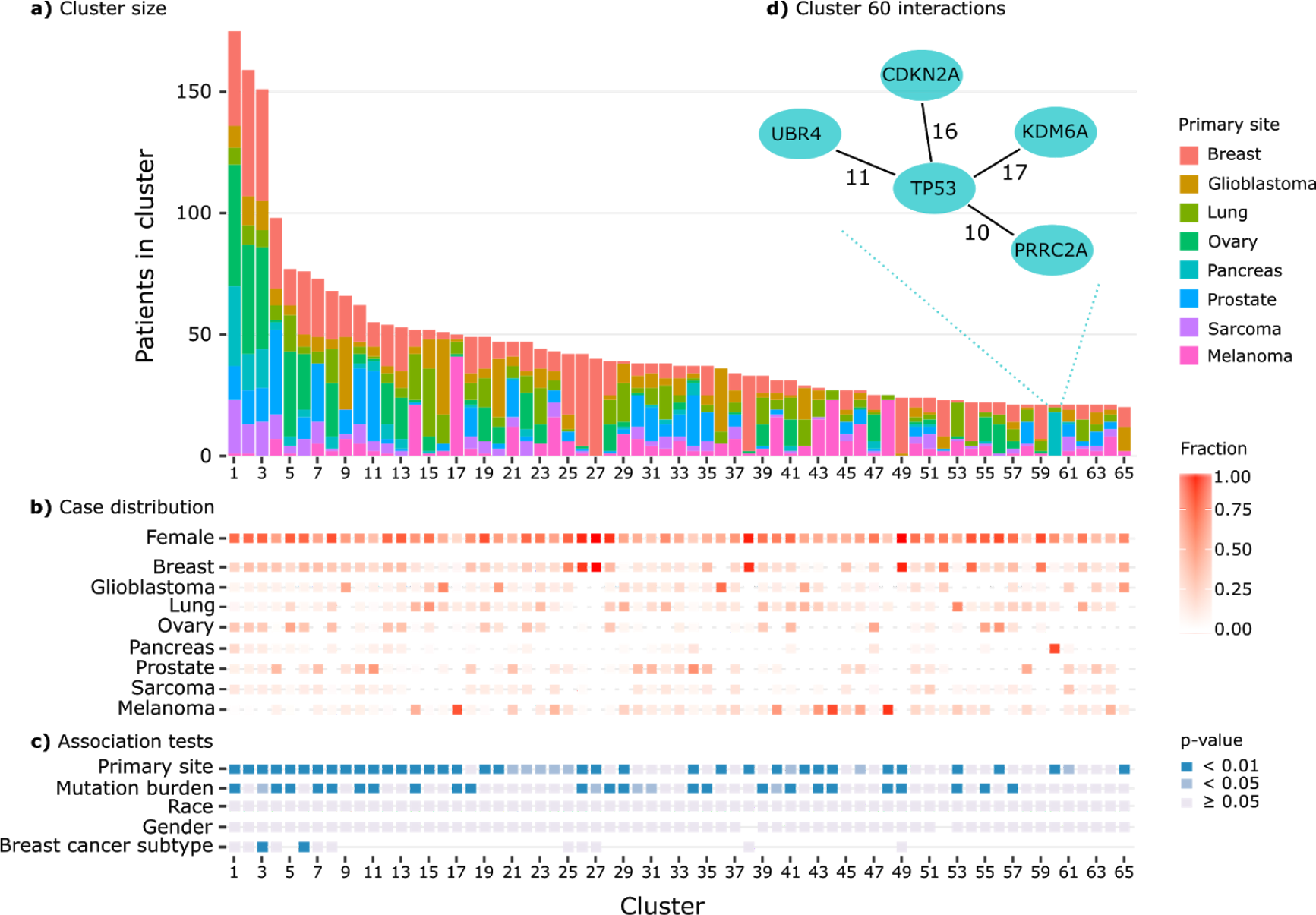
Distribution of EPICC clusters in the TCGA 8-cancer dataset. EPICC clusters varied widely in number of patients in the cluster and primary site distribution (a, b). Clusters were often biased in their primary site composition and some were associated with cancer mutation burden, but no gender or race bias was observed (c). The major interactions driving cluster 60 are shown to illustrate the EPICC score (d); ellipses represent interacting proteins, and lines are labelled with the number of sample pairs with co-mutation in the connected proteins. EPICC clusters with at least 20 patients only are shown; p-values are corrected for familywise error rate (Holm’s step-up procedure) considering 271 tests. In (c) the absence of a box indicates the relevant test was not performed due to structural zeroes or low pre-test power.

Examination of the shared mutations in selected clusters suggested that the EPICC clustering was reflective of known biology. 73% of cluster 36 consisted of glioblastoma patients, and co-mutation of the classic interacting drivers *PTEN* and *EGFR* was a notable feature in this cluster (Crespo *et al.*, 2015). Cluster 60 was strongly associated with pancreatic cancer. Interestingly, this pancreatic cancer cluster was not driven by mutations in *KRAS* or *TP53*, the dominant genes affected in pancreatic cancer (Biankin *et al.*, 2012), but rather by co-mutation of second-tier genes *CDKN2A, KDM6A, PRRC2A*, and *UBR4*, which through their mutual interactions with p53 led to a high EPICC similarity score between patients in the cluster (Figure 2d). This behaviour underscores the gain in sensitivity that can be achieved using a network-based clustering like EPICC.

Intriguingly, some EPICC clusters correlated with cancer subtype, rather than primary type. Clusters 3 and 6, although not strongly linked to breast cancer as a whole, were significantly associated with the triple negative and HER2 positive subtypes of breast cancer (both p < 0.003 after Holm’s correction for 271 tests, Figure 2c). Investigation of mutated genes in these clusters revealed *TP53* mutation as a central driver for membership in these triple negative / HER2 clusters, consistent with known features of these breast cancer subtypes

(Darb-Esfahani *et al.*, 2016). In contrast, clusters 25, 26, 27, 38, and 49, which were associated with breast cancer but not biased towards a specific subtype, were characterised by co-mutation of *CDH1*, *MAP3K1*, and *PIK3CA*, and seldom involved *TP53* changes.

Motivated by the observation that although some EPICC clusters were reflective of known cancer mechanisms, many clusters did not have a clear biological annotation, we undertook a survey to identify protein modules inside EPICC clusters which may be indicative of novel cancer biology.

### Identifying cancer cliques inside EPICC clusters

We developed the concept of a “cancer clique” to investigate patterns of recurrent mutation inside EPICC clusters. Cancer cliques are sets of fully interacting proteins which are disrupted in every patient inside a cluster (Figure 1c,d). Such fully interacting sets of proteins are likely tightly-coupled multi-protein complexes, for which the disruption of any single member protein would affect function of the entire complex. Thus, cancer cliques represent an estimate of disrupted functional modules or biological mechanisms within an EPICC cluster.

We observed that most EPICC clusters contained several cancer cliques, and that no single clique can represent the shared similarity of all patients within one cluster. Based on this finding, we identified 48 cliques that were present in several clusters, that might represent important recurrently disrupted biological mechanisms. To focus on novel biology, we filtered out cliques that aligned with known pathways, resulting in 28 novel cliques (Figure 3).

**Figure 3:**
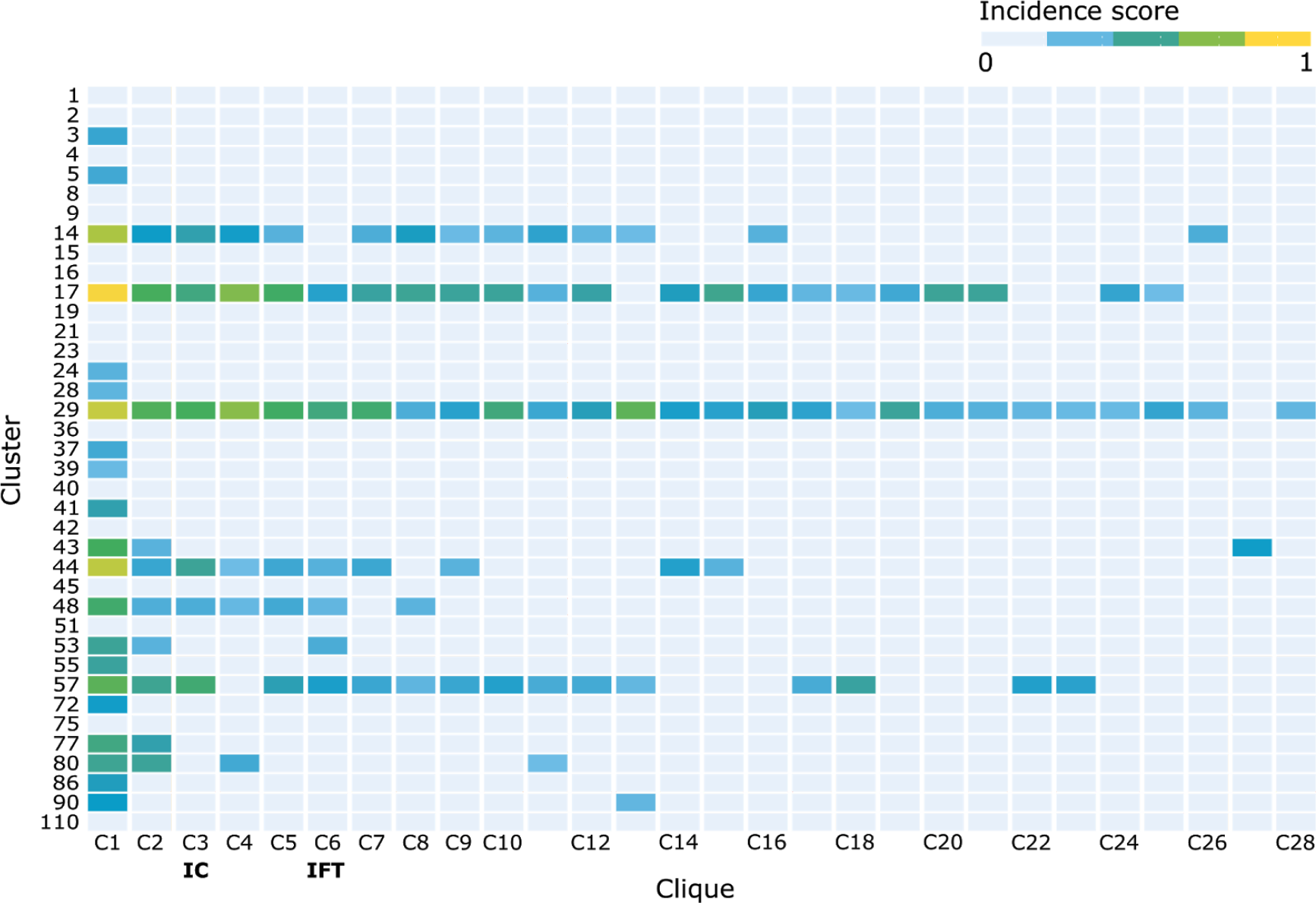
Recurrent novel cancer cliques. After excluding cliques with known function, 28 novel cliques (columns) were identified across all clusters. These cliques were present in 38 EPICC clusters (rows). Cells denote the composite score describing the incidence of a given clique in the row cluster, with higher values indicating that patients in the row cluster are highly enriched for mutations in the column clique. Two cliques were selected for further characterisation, the Integrator Complex (IC) and the Intraflagellar Transport group (IFT).

We selected two novel cliques for illustration based on preliminary association with patient outcome in the TCGA cohorts. Clique C3 contained all 15 named members of the Integrator Complex (IC), and thus we termed this the Integrator Complex clique. Similarly, clique C6 contained predominantly proteins involved in intraflagellar transport; this clique we also termed the Intraflagellar Transport (IFT) clique. Interestingly, although these cliques could be linked to likely biological processes manually, they were absent from the MSigDB pathways database.

### EPICC reveals cliques not identified by pathway analysis

Of the 48 robust recurrent cancer cliques identified inside EPICC clusters, 20 (42%) had significant overlap with known signalling pathways described by the MSigDB database (Subramanian *et al.*, 2005; Liberzon *et al.*, 2011), corroborative of biological meaning embedded in these cliques. Notably, the remaining 28 cancer cliques had no significant correlate in MSigDB, and thus could not have been identified by a MSigDB pathway-based analysis. MSigDB is a comprehensive and representative pathway database, and this result underscores the value and increased sensitivity from undertaking a global unbiased approach to module discovery, as exemplified by EPICC.

### The EPICC resource

We have developed an interactive front-end for the investigation of EPICC clusters and cliques identified in the eight-cancer TCGA cohorts, displaying network context and correlations between clique mutation and outcome. This resource is freely-available for researchers wishing to explore EPICC clusterings, at https://github.com/shu2010/EPICC.

## Discussion

Here we have described EPICC, a novel approach to cluster cancers based on mutated genes. EPICC overlays mutation data on the protein-protein interaction network, to provide a unique prism through which to examine cancer subtypes. EPICC clusters can be further interrogated for constituent cancer cliques, which may reflect novel cancer processes that are not captured by methods dependent on protein annotation. The EPICC methodology is broad in its applicability and amenable to a range of refinements, which may represent fruitful directions for future development. EPICC has been implemented on the Vodafone DreamLab platform and applied to a set of cancers from the TCGA project, and we make our results available to cancer researchers interested in interrogating the TCGA data for cancer cliques.

EPICC both recapitulated known cancer groupings, and revealed potentially significant cross-cancer clusters (Figure 2a-c). Notably, EPICC identified cancer-consistent clusters even when the dominant mutations were not in interacting proteins; for example in pancreatic adenocarcinoma the majority of cancers bear mutations in *KRAS* and *TP53* (Biankin *et al.*, 2012), but these proteins do not interact and so can not directly contribute to EPICC clustering. Despite this, EPICC clustered pancreatic cancers together on the basis of lower-frequency shared mutations, demonstrating the robustness of the method. Such robustness is an attractive feature of transcriptomic clustering approaches, with DNA-based methods considered more fragile (Ray *et al.*, 2014). EPICC’s ability to leverage protein interaction network information to extract robust clusters from DNA mutations is a distinct advantage, particularly in clinical contexts where transcriptomic measurements are challenging to acquire.

Our interrogation of EPICC clusters identified a number of cancer cliques which may represent functional modules that are commonly disrupted in cancer. More than half of the cancer cliques we identified did not map to known pathways in the comprehensive MSigDB database, underscoring the advantage of a network-based approach to cancer clustering and module identification that is not reliant on potentially biased databases.

Two of the novel cliques we identified could be linked to known biological modules with roles in cancer: the Integrator Complex (IC) and the Intraflagellar Transport (IFT) proteins. The IFT clique includes 13 proteins involved information and maintenance of the primary cilium, which is integral to a number of cancer signalling pathways (Taschner and Lorentzen, 2016). The IC clique contains 15 members of the Integrator Complex, a major component of the RNA polymerase II mediated transcription machinery (Rienzo and Casamassimi, 2016). The Integrator Complex is believed to mediate post-transcriptional control of developmental genes and transduction cascades in response to stress, growth factors or other stimuli (Gaertner and Zeitlinger, 2014), and has been identified as a common target for mutation in cancer (Federico *et al.*, 2017). The correlation between these cliques and biological modules linked to cancer processes suggests that other novel cancer cliques will be fertile ground for further investigation.

Our illustration of EPICC on TCGA cancer data represents a first demonstration of the technique, and a number of refinements are being explored. In this work we considered all non-synonymous somatic variants as equivalent, and a variant of EPICC that weights variants by predicted effect on their protein will likely reduce noise, particularly in the case of high mutational burden cancers such as melanoma (Chalmers *et al.*, 2017). Numerous adjustments to the protein network are also possible, such as more stringent curation of interactions, longer range or weighted interaction scoring, or the consideration of directed interactions (Vinayagam *et al.*, 2011). Finally, we did not exhaustively explore clustering thresholds or cancer clique cutoffs, parameters which may further improve the purity of detected clusters and cliques.

EPICC resulted from the first projects to be completed on the Vodafone DreamLab platform, Project Decode and Genetic Profile. The DreamLab platform provided a unique computational resource that was well-suited to the highly distributed calculation of the EPICC metric, but may be more limited for other applications. In our experience the DreamLab platform is particularly appropriate for extremely parallel tasks that involve high-intensity calculations on small data packets, where rapid analysis iteration is not required and input data and analysis code are stable over many months.

## Methods

### Source Data

#### Protein interaction network

EPICC scores were computed using the BioGRID protein-protein interaction (PPI) network Release 3.4.160 (Chatr-Aryamontri *et al.*, 2017). A protein pair was considered to interact if it contained any of the following BioGRID physical protein-protein interactions: Affinity Capture-Luminescence, Affinity Capture-MS, Affinity Capture-Western, Co-crystal structure, Co-fractionation, Co-purification, Far-Western, FRET, Protein-Fragment Complementation Assay, Protein-peptide, Proximity Label-MS, Reconstituted Complex, and Two-hybrid. Interactions due to biochemical activity, such as phosphorylation or ubiquitination, were excluded.

#### Somatic mutations

3,750 cancers were identified in The Cancer Genome Atlas (TCGA), constituting 985 breast invasive carcinomas, 566 lung adenocarcinomas, 493 prostate adenocarcinomas, 436 ovarian carcinomas, 467 skin melanomas, 391 glioblastomas, 237 sarcomas, and 175 pancreatic ductal adenocarcinomas. Somatic mutations as of 2017-12-02 were extracted for these cancers using the R Package TCGABiolinks (Colaprico *et al.*, 2016). A protein was considered mutated in a sample if it was affected by at least one non-silent somatic mutation (consequence fields in the set Frame_Shift_Del, Frame_Shift_Ins, In_Frame_Ins, In_Frame_Del, Missense_Mutation, NonSenseMutation, Splice_Site, Translation_Start_Site, and Nonstop_Mutation), as called by at least one variant caller in the set VarScan, MuSE, MuTect, and Somatic Sniper.

### EPICC Scores

#### Raw distance calculation

Raw undirected EPICC similarity scores between all *n* = 3750 cancers were computed as the matrix product *C* = *M^T^ AM*, where *C* is a 3750 × 3750 matrix of raw EPICC scores, *A* is a 20871 × 20871 adjacency matrix describing the protein interaction network (*A*_*i,j*_ = 1 if an interaction is observed between proteins *i* and *j*, else *A*_*i,j*_ = 0), and *M* is a 20871 × 3750 matrix of sample mutation indicators (*M*_*i,k*_ = 1 if at least one non-silent mutation in protein *i* is observed in sample *k*, else *M*_*k,i*_ = 0). The raw EPICC scores for all unique pairwise sample combinations can be obtained from either the upper or lower off-diagonal elements of matrix *C*. Although not explored in this work, we note that EPICC similarity scores can also be calculated for directed graphs by a modified procedure, which may be advantageous in well-annotated contexts such as kinase pathways (Vinayagam *et al.*, 2011).

#### Normalisation

Raw EPICC scores have a variable scale, dependent on the mutation frequency and mutated proteins in a given sample pair. To enable comparison of EPICC score magnitude across sample pairs, we undertook quantile normalisation of scores against an empirical null. We generated 1,000 random protein-protein interaction networks using a procedure that conserved interaction degree between the original and randomised networks (Molloy and Reed, 1995). For each patient pair, EPICC scores were calculated on each of the randomised networks, yielding a pair-specific null distribution of EPICC scores. The observed raw EPICC score for a patient pair was then quantile normalised against this empirical null for that same pair, to produce a normalised EPICC score. Network generation was performed using the igraph R package (Kolaczyk and Csárdi, 2014), and null scores were computed on the Vodafone DreamLab platform.

### Clustering

Given normalised EPICC similarity scores between each pair of patients, we clustered patients by affinity propagation (Frey and Dueck, 2007), as implemented in the R package apcluster (Bodenhofer *et al.*, 2011). The square of the normalised EPICC score was used as the clustering similarity metric, and the median pairwise similarity metric as the input preference parameter. All other parameters were left at defaults.

All patient clusters containing more than 20 patients were tested for association with primary site, race, gender, and mutation burden. Additionally, patient clusters containing more than 20 breast cancer cases were tested for association with breast cancer subtype, and clusters containing more than 20 lung adenocarcinoma cases with pack years smoked. Tests against discrete variables were performed using Fisher’s exact test as implemented in R, for each cluster comparing class frequencies within the cluster against frequencies for all samples outside the cluster. Fisher’s test p-values were computed exactly for 2 × 2 comparisons, and approximated with 10^5^ permutations for larger tables. Mann-Whitney tests were used to compare mutation burden and pack years between samples within a group to those outside the group. In total 271 tests were performed; p-values are reported following familywise error rate control using Holm’s step-up procedure (Holm, 1979). All clinico-pathological variables were sourced from TCGA data tables; breast cancer subtype was derived from reported ER, PR, and HER2 IHC staining positivity. Mutation burden for a sample was the count of proteins bearing non-synonymous mutations.

### Cancer Clique Discovery

The mutations corresponding to each cluster were transformed into PPI subgraphs and interrogated for prevalence of complete polygonal subgraphs or cliques. All possible maximal cliques were identified using a variant of the Bron–Kerbosch algorithm implemented in R package igraph (Kolaczyk and Csárdi, 2014). Only clique(s) that persisted after removal of promiscuous proteins (degree ≥ 100) from the network were retained as high confidence cliques.

#### Identification of recurrent novel cliques

Robust cliques recurrent across EPICC clusters were selected by a multi-stage filtering procedure. First, highly stable EPICC clusters were identified using five affinity propagation clustering runs, using exemplar preferences from the 10th to the 50th percentile of similarities. Clusters identified in at least 4 of 5 runs were retained, and cliques extracted. As some redundancy was observed between cliques, redundant cliques were merged by Jaccard similarity (threshold > 0.8), and especially small (fewer than five proteins), or large (more than 30 proteins) cliques were excluded, as they were either too small for pathway correlation, or too large for specific activity annotation. A total of 48 cancer cliques derived from 38 EPICC clusters remained following this process. Cliques were tested for significant overlap with MSigDB signalling pathways (Liberzon *et al.*, 2011; Subramanian *et al.*, 2005), and a function assigned if a significant association was detected (Holm-adjusted p-value < 0.05, hypergeometric test). 28/48 cliques had no significant pathway association, and were ranked by the product of mutated patient ratio and percentage of mutated clique elements in every cluster.

